# Overview of the expression of key regeneration genes in embryo development and matured tissues in Axolotl (*Ambystoma mexicanum*)

**DOI:** 10.1101/2020.09.24.312462

**Authors:** Harshil Shah, Juan Caballero

**Affiliations:** SkoolMentor; Universidad Autonoma de Queretaro

**Keywords:** Axolotl, tissue regeneration, expression analysis, RNA-seq

## Abstract

The axolotl is a Mexican endangered species with the capability to perform tissue regeneration in amputated extremities. Being an important research model, the complete genome of the Axolotl was sequenced in 2018 for the first time, revealing an enormous genome: the largest of any animal ever sequenced, and about 10 times larger than the human genome, a new landmark achievement in biology research.

In this report, we collected 70 known genes that play an important role in tissue regeneration and searched for their expression during embryo development, regeneration, and in 6 adult tissues: the heart, liver, gills, front leg, rear leg, and tail. We observe those 70 genes expression levels in the 3 conditions and approximately 3 genes seem to be expressed exclusively in regeneration.

This report displays a few insights on how this marvelous species can modulate its regenerative capabilities. There is still much more to explore with the Axolotl. Further research into their regenerative capabilities could provide researchers with the perspicacity to possibly accomplish human limb regeneration.

## INTRODUCTION

The axolotl (*Ambystoma mexicanum*) is famous for its lifelong youthfulness, more scientifically described as neoteny, the fact that it does not undergo the metamorphosis from larval, aquatic stage to adult, terrestrial salamander-like related amphibian species would, but reaches maturity while in the larval form (Gross 2018). The Axolotl is also paedomorphic, retaining juvenile features throughout their lifespan and into adulthood, including the external gills, a body fin, and moveable eyelids (https://animals.sandiegozoo.org/animals/axolotl).

Diverse molecular analysis has been performed on this species, including the complete genome of the Axolotl which was sequenced in 2018 for the first time. The genome revealed enormous chromosomes: it is the largest of any animal ever sequenced, and about 10 times larger than the human genome (Dockrill 2020). The full sequencing marked a landmark in Biology because the data available could be researched for potential awareness into their regenerative ability as well as other useful traits the Axolotl possesses.

The axolotl’s capability allows them to regenerate their limbs, lungs, heart, jaws, spines, and even parts of their brain (https://animals.sandiegozoo.org/animals/axolotl). Studies have found that they can regrow a new limb five times perfectly in a few weeks. Other than being regenerative, wild Axolotl’s can change the hue of their skin a few shades for camouflage.

The stages of regeneration for a salamander leg are visible in the composite images Figure 1, Figure 2, and Figure 3. The intact limb is at the left; the series to the right shows the gradual regeneration of the limb over a couple of months. The steps include: first the intact limb receives trauma, and we see the wound without the limb.

**Figure 1.**
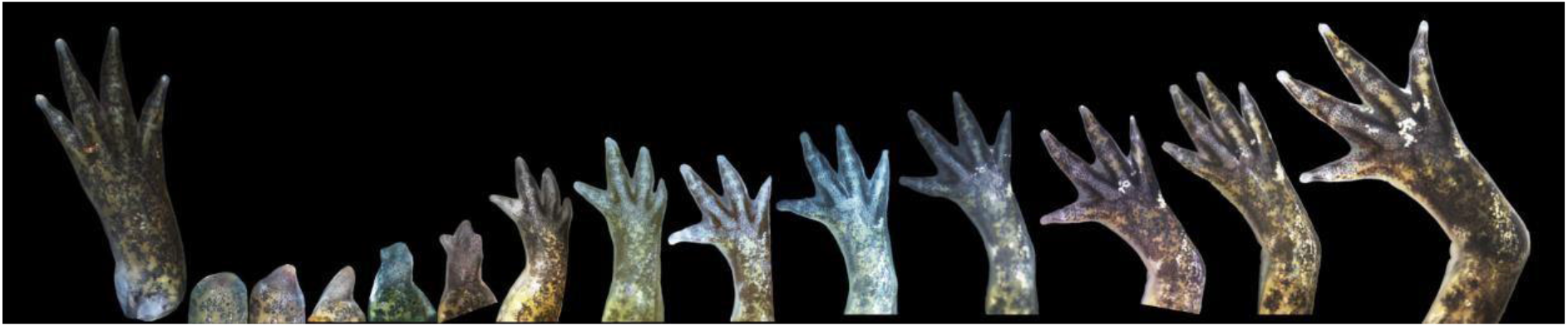
Photographs of limb regeneration in Axolotl (Farrow 2017)

**Figure 2.**
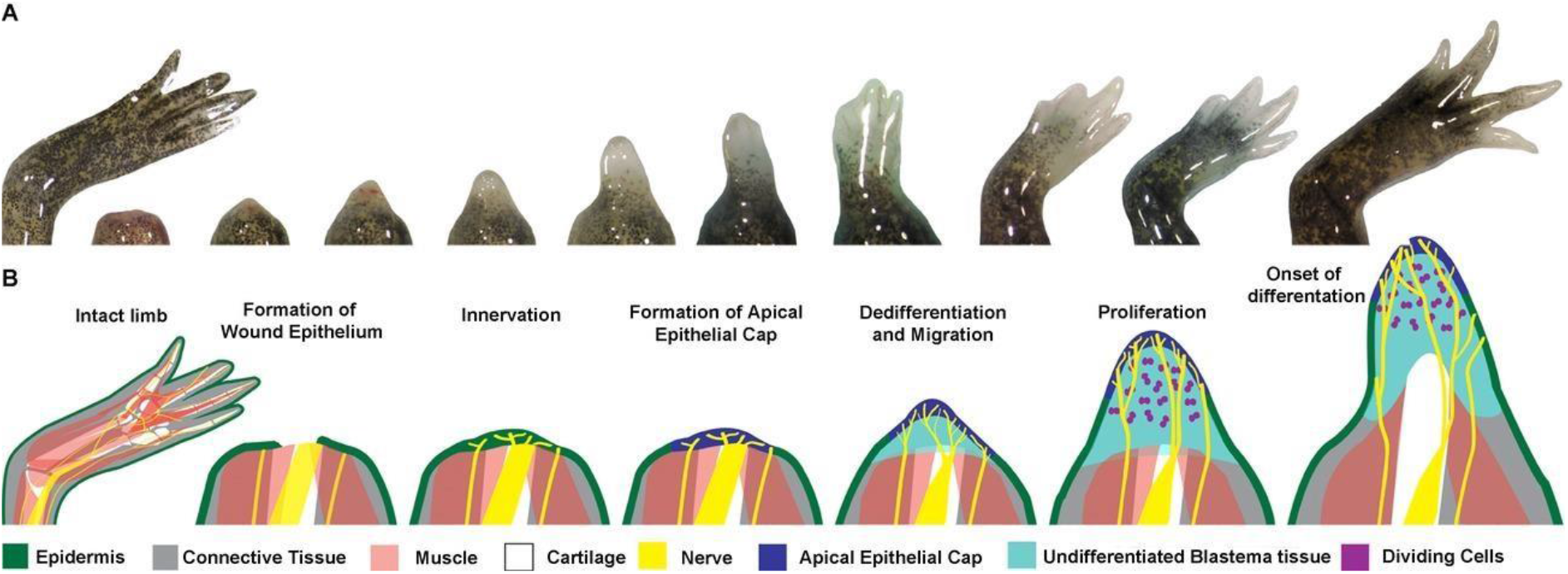
Graphic representation of the limb regeneration in Axolotl (McCusker 2015)

**Figure 3.**
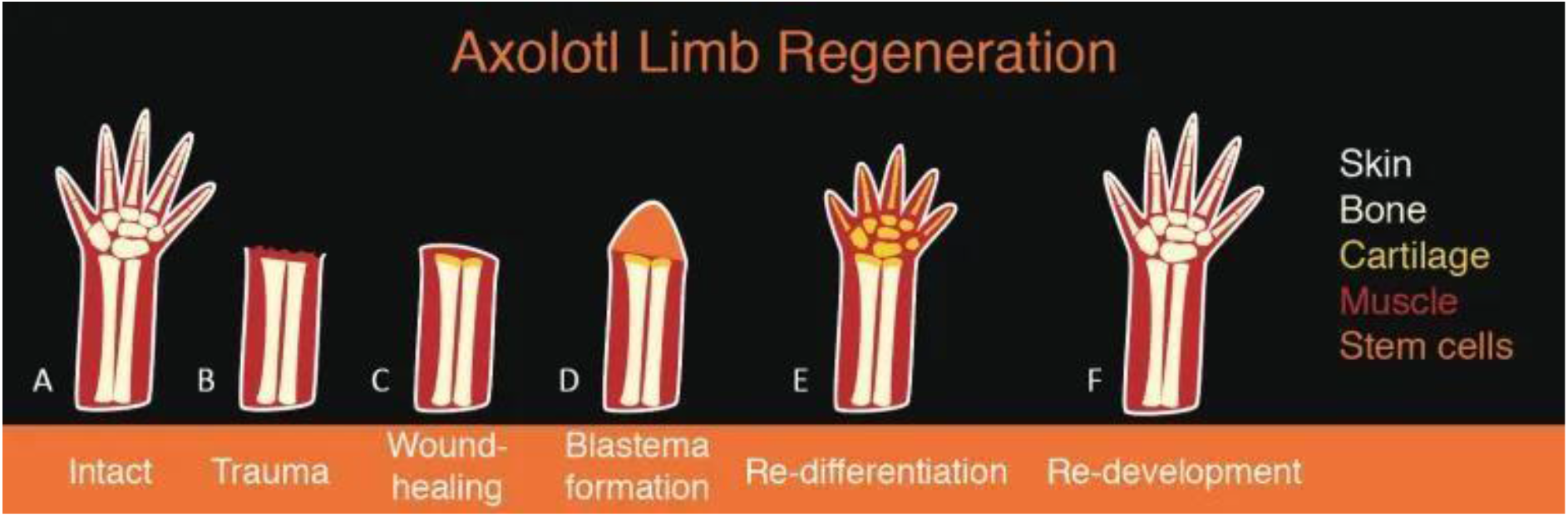
Bone and muscle regeneration in Axolotl limb amputation (Dunlap 2018)

The first step of regeneration involves epithelialization, a process of covering excoriated or traumatized epithelial surfaces. First, a clot of blood cells rapidly stops bleeding at the wounded site. Then, a layer of cells begins to form an epithelial and cover the wound completely (Dunlap 2018). The cellular and molecular processes involved in initiation, maintenance, and completion of epithelialization are essential for successful wound closure.

Next, in innervation, nerves are replaced into the wounded areas and go into muscle fibers, innervating the muscle fiber. The wounded epidermis transforms into a layer of signaling cells called the Apical Epithelial Cap, which has a vital role in regeneration. Interesting research conducted previously further analyzes how cells at the stump of the wound do not revert to a completely embryonic state as previously thought, but rather the parts that were muscle remembers that it needs to grow muscle, whereas the part that was nerve remembers that it needs to grow nerve. They are still able to grow into tissues, but only certain kinds of tissue (Farrow 2017).

The cells underneath the epidermis also begin to rapidly divide, forming a cone-shaped structure known as a blastema. The cells that make up the blastema are thought to be bone, cartilage, muscle, or other cells that de-differentiate to become like stem cells.

Dedifferentiation is an important biological phenomenon whereby cells regress from a specialized function to a simpler state reminiscent of stem cells. Stem cells are self-renewing cells capable of giving rise to differentiated cells when supplied with the appropriate factors, and these stem cells can then begin migrating to necessary areas.

These de-differentiated stem cells in the blastema then grow and multiply, eventually regaining their identity as fully developed cells to form necessary tissues and organs. As the blastema and its cells continue to divide, the growing structure flattens and eventually resembles a perfect copy of the lost limb, including nerves and blood vessels that are connected to the rest of the body.

Further research from James Godwin, the lead study author, from the Australian Regenerative Medicine Institute (ARMI) at Monash University in Melbourne, concludes that macrophages play an essential role in the regenerative process. They experimented with salamanders by injecting the animals with a chemical substance that depleted the macrophage-levels of the animals and then amputating limbs to see the body’s reaction. They found that those salamanders could not regenerate the limbs and had severe scarring build-up. However, after they reverse-engineered this and re-introduced the macrophages, the animal was able to regenerate those same lost limbs, thus demonstrating the importance of macrophages in the process (Lewis 2013). Macrophages have already been known to play crucial roles in mouse embryo tissues and development for many animals. Godwin believes that the ability to regenerate was turned off due to evolution, but it may be possible to reevaluate that and reverse the process (Lewis 2013).

### Axolotl genome and research

Learning more about the Axolotl’s anatomy, geographic density, and lifestyle/cycle could contribute to the overall understanding of the animal’s abilities and tendencies. Studying the major organs of the Axolotl could provide information on the roles of different genes In molecular processes and metabolism. Since our research and experiment focused on the Axolotl’s Rear and Front Leg, Gills, Heart, Liver, and Tail, Figure 4 shows a diagram of the Axolotl body labeled with major organs.

**Figure 4.**
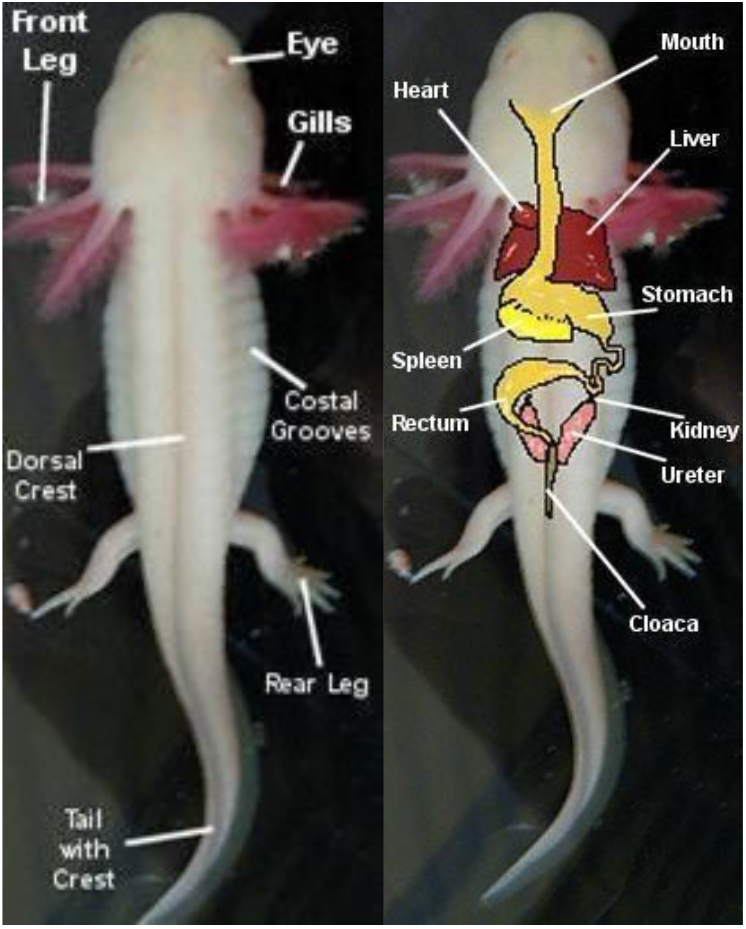
Axolotl main organs.

The axolotl’s capability allows them to regenerate their limbs, lungs, heart, jaws, spines, and even parts of their brain (https://animals.sandiegozoo.org/animals/axolotl). Studies have found that they can regrow a new limb five times perfectly in a few weeks. Other than being regenerative, wild Axolotl’s can change the hue of their skin a few shades for camouflage.

The final superpower is that the Axolotl is 1,000 times more resistant to cancers than mammals, and researchers from the University of Nottingham used an extract from axolotl eggs to stimulate tumor suppressor genes and prevent breast cancer growth (Travers 2018). Scientists from the University brought the Breast Cancer Cells back under control by reactivating their tumor suppressor genes (University of Nottingham 2011). Treating the cells with Axolotl oocyte extract reactivated the suppressor genes and stopped the cancer from growing, and within 60 days, there was no cancerous growth. Cancers are caused by mutations within the cell division cycle and uncontrollable growth of mutated cells, and among the most important of these genes are tumor suppressor genes. These genes repress the development of cancers and normally act as a control point in the cell division cycle. Therefore, the switching off tumor suppressor genes is a common cause of cancers, including breast cancer” (University of Nottingham 2011). Using novel technology that makes use of the eggs of the axolotl salamander, cancer cells were treated with extracts made from axolotl oocytes that could reverse the epigenetic marks on tumor suppressor genes, causing these genes to reactivate, and thereby stopping the cancerous cell growth. Axolotl oocytes are packed with molecules that have very powerful epigenetic modifying activity. Previously, the Johnson’s lab at the University of Nottingham showed that extracts prepared from these oocytes have a powerful capacity to change epigenetic marks on the DNA of human cells (University of Nottingham 2011).

For this research, the three major functions we focused on were limb regeneration, development in the embryo, and major organs, as mentioned before. Focusing on each, first, we took data from experiments when Axolotls were amputated limbs in a laboratory and studied over a 40-period study. The key stages of regeneration, as the data reflected, were the: 1, 2, 3, 4, 5, 6, 7, 8, 9, 10, 11, 12, 14, 16, 19, 24, and 40 stages. For the embryo development, Figure 5 shows the 40 stages of development of the Axolotl from an egg to larva. Finally, we included data of the expression levels of proteins in the Axolotl in major organs, as listed above: Front and Rear Leg, Tail, Gills, Heart, and Liver.

**Figure 5.**
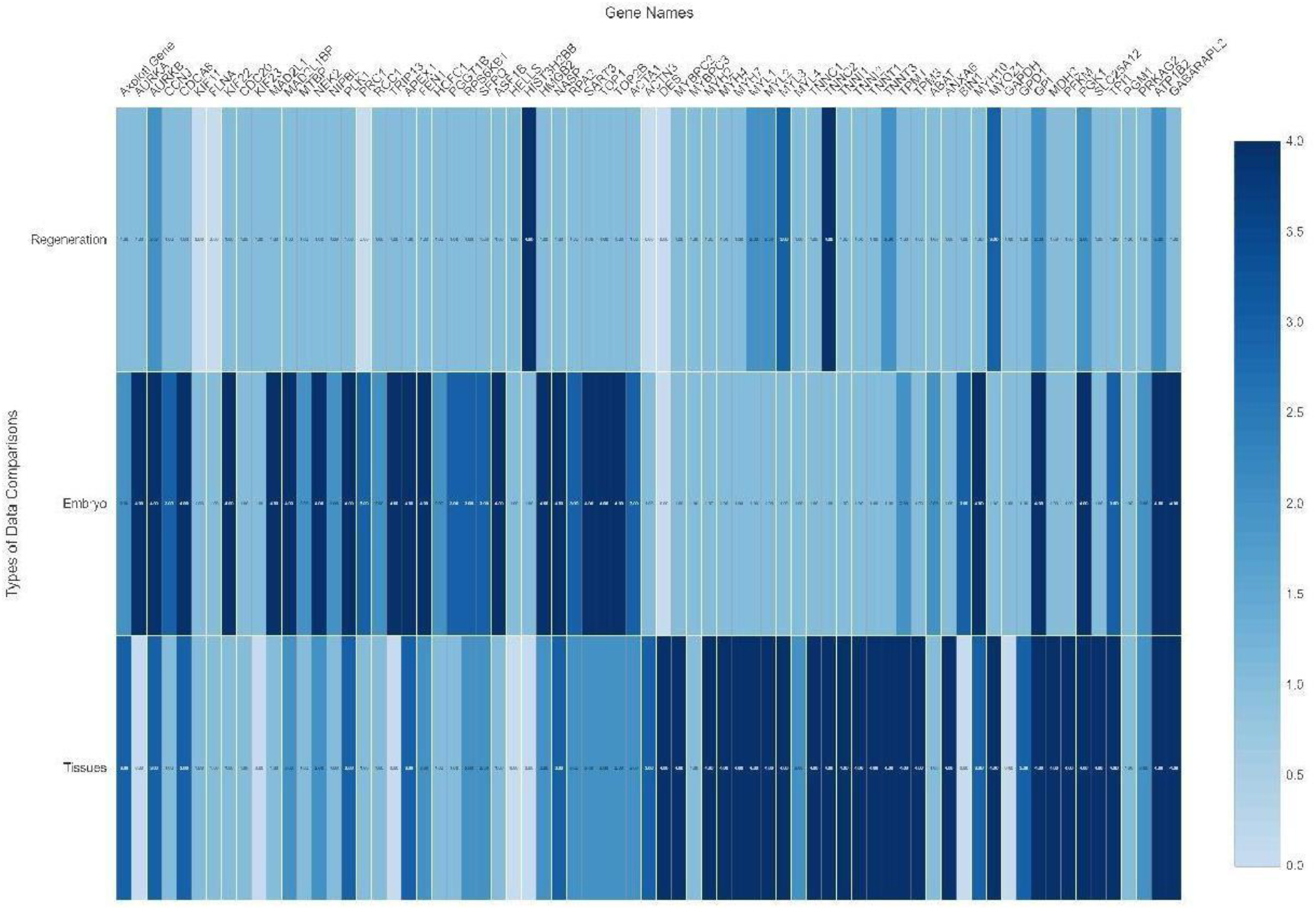
Heatmap for gene expression in the 3 datasets.

## METHODS

### Sequences

We used the sequences of all Axolotl assembled transcriptome reported in (Caballero-Perez 2018), in brief, such database was built using the assembly combination of regeneration transcriptome (Bryant 2017), embryo development (Jiang 2017) and the RNA-seq of the adult tissues. After merging the assemblies, the transcriptome was quantified for each stage with Kallisto and normalized in transcript per million to enable cross-comparisons.

### Selection of key genes in regeneration

We used the list of 70 key genes involved in Axolotl regeneration as reported in (Sibai 2019), in the initial screening we noted many genes were incomplete, therefore we collected the orthologue gene searching the main human RefSeq protein for each gene in NCBI GenBank.

### Transcript identification in Axolotl transcriptome

We utilized NCBI Blast+ to search the transcriptome database using “tblastn” to use the human protein as a query, and the transcriptome nucleotide database, so all transcripts are translated to the equivalent proteins for the comparison. With the specific Axolotl genes that were most similar to the human gene found (E-value < 1e-6), we collected those transcripts identifiers and searched the database for their expression levels in the three major functions: regeneration, development, and tissues.

## RESULTS

### Expression patterns

All 70 genes were identified in Axolotl transcriptome. The genes were classified into different levels of expression based on the greatest number of stages/time-periods in which the transcript was expressed at that level. No expression was considered for 0 tpm, low expression 0-2 tpm, medium expression 2-5 tpm, high expression 5-20 tpm, and extreme high expression was >20 tpm.

As shown in Table 1, in regeneration, most genes had low expression while a few had high and extremely high expression. The few genes that are left are expressed in extremely high quantities. In the embryo/development, more genes were expressed in extremely high quantities as well as many genes that were expressed in lower quantities. Finally, in the major organs of the Axolotl, most genes were extremely high expressed out of all the three functions

**Table 1.**
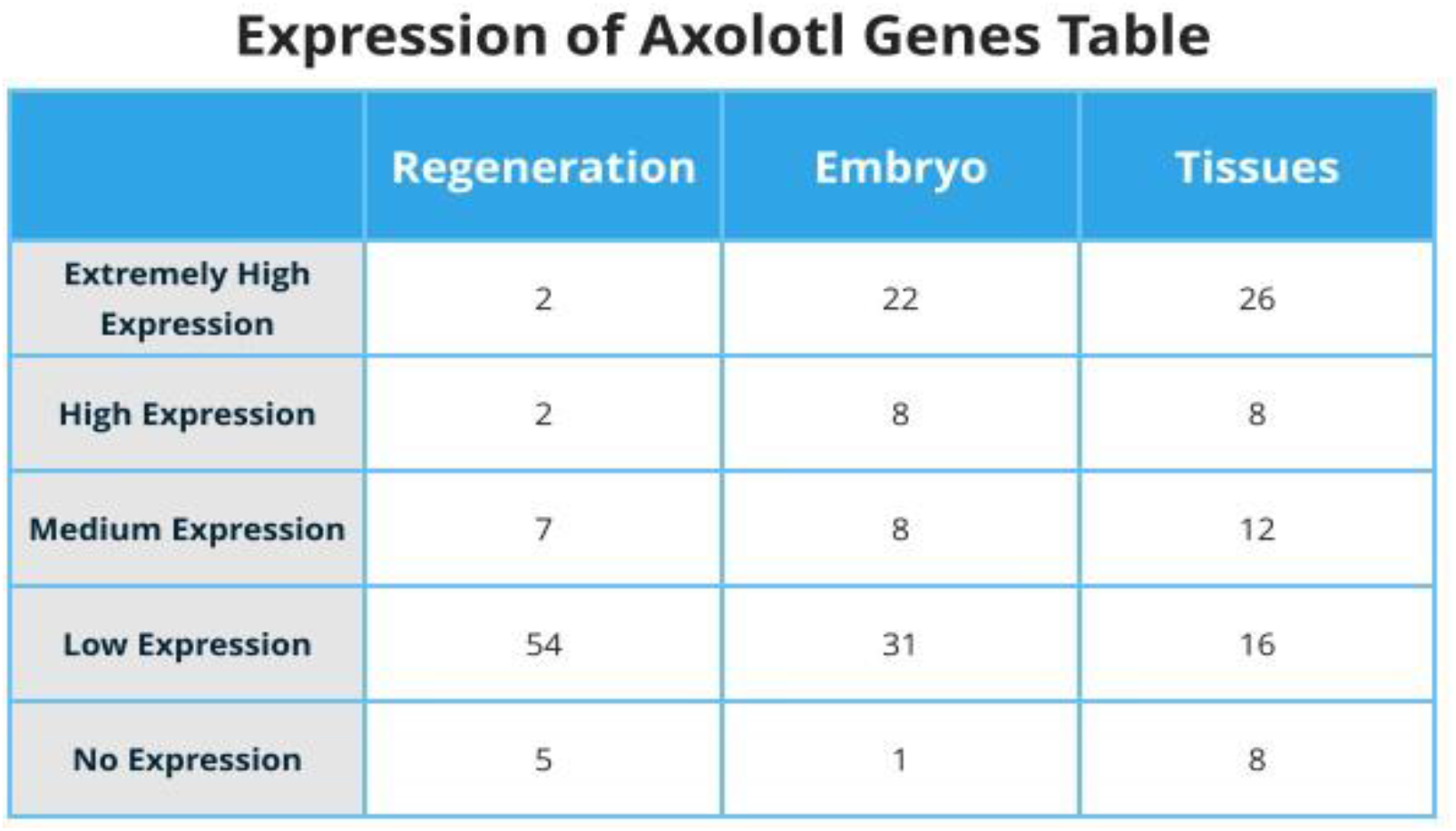
Global expression of genes in the 3 transcriptomes.

**Expression of Axolotl Genes Table**
|  | Regeneration | Embryo | Tissues |
| --- | --- | --- | --- |
| Extremely High Expression | 2 | 22 | 26 |
| High Expression | 2 | 8 | 8 |
| Medium Expression | 7 | 8 | 12 |
| Low Expression | 54 | 31 | 16 |
| No Expression | 5 | 1 | 8 |

Table 2 displays the expression for each of the genes. The first column lists the gene symbol that was being inputted into the transcriptome database. The next column lists the Axolotl gene identifier that was shown to be most like the human gene, and it is listed as in the Amex transcriptome database. The next three columns list the expression levels of the Axolotl genes in regeneration, development, and major organ tissues in detail, with all the different expression levels present throughout the experiment.

**Table 2.** Detailed gene expression in the 3 main datasets, expression levels are classified as: Low expression: 0 - 2 tpm, Medium expression: 2-5 tpm, High expression: 5-20 tpm, and Extremely High: >20 tpm.

| Gene Symbol | Axolotl Gene | Expression in Regeneration | Expression in Embryo / Development | Expression in Axolotl Tissues |
| --- | --- | --- | --- | --- |
| AURKA | Amex3_3159454 | Low expression throughout process | Extremely high expression, gradually decreasing | No expression |
| AURKB | Amex3_3260416 | Medium expression in regeneration, increasing gradually towards later stages | Very high expression, gradually decreasing | High expression in tissues (Especially front and rear legs, gills, and tail) |
| CCNJ | Amex3_3258442 | Low expression | High expression, decreases substantially at end | Low expression 1 |
| CDC20 | Amex3_3534199 | Low expression in the 3rd, 4th, and 7th stage (6hrs, 12hrs, 5d respectively) | Low expression, with medium expression in the 7th and 11th stage | Low expression in the heart, rear leg, and tail |
| CDCA8 | Amex3_2115615 | Low expression | Extremely high expression in all stages of development | High expression (14+ in Front and Rear legs, Tail, and Gills) |
| KIF11 | Amex3_208660 | No expression | Low expression only in the 1st, 2nd, 9th, 12th, 14th, 16th, and 19th stages | Low expression only in Front leg and tail |
| FLNA | Amex3_208660 | No expression | Low expression only in the 1, 2, 9, 12, 14, 16, 19, and 24 stages | Low expression only in Front leg and Tail |
| KIF22 | Amex3_3258744 | Low expression | Extremely high expression, gradually declining | High expression in Gills and Rear leg, Low expression in rest |
| KIF23 | Amex3_2629094 | Low expression | Low expression in only 2, 4, 9, 12, 16, 19, 24, 40 stages | No expression |
| MAD2L1 | Amex3_3532025 | Low expression | Extremely high expression, decreases to medium expression after 11 stage | Low expression |
| MAD2L1BP | Amex3_3533474 | Low expression | Extremely high expression, decreases to medium expression after 24 stage | Medium expression |
| MTBP | Amex3_3531074 | Low expression at 12hrs, 3d, 10d, 21d, 28d | Medium expression in 1, 2, 3, 4 stages, low expression in all other stages | Low expression |
| NEK2 | Amex3_3259832 | Low expression | Extremely high expression | Low expression in Heart and Liver, Medium expression in the rest |
| NIPBL | Amex3_208253 | Low expression only at 6hrs, 1d, 14d, 21d, 28d | Medium expression in middle stages, Low expression in rest | Low expression |
| PLK1 | Amex3_402983 | Low expression | Extremely high expression, decreasing gradually | Low expression in Heart and Liver, High expression in rest |
| PRC1 | Amex3_1857349 | None | High expression in the first and last stages, extremely high expression in the middle | Low expression |
| RCC1 | Amex3_819365 | Low expression 10d, 21d | Medium expression | Low expression in only Front leg |
| TRIP13 | Amex3_3159454 | Low expression at 3hrs, 6hrs, 1d, 3d, 5d, 14d, and 28d | Extremely high expression, decreases substantially at end to low expression | No expression |
| APEX1 | Amex3_2503633 | Low expression | Extremely high expression | High expression |
| FEN1 | Amex3_3533277 | Low expression | Medium to High expression | High expression in tail, medium expression in rest |
| HCFC1 | Amex3_2112644 | Low expression | Low to Medium expression | Low expression |
| PGGT1B | Amex3_3156215 | Low expression | Medium expression in first two stages, then high expression | Low expression only in Front and Rear leg, Gills, and Heart |
| RPS6KB1 | Amex3_3106233 | Low expression | Low expression in first two stages, medium expression in next three stages, then high expression | Medium expression |
| SFPQ | Amex3_1628 | Low expression at 0hrs, 12hrs, 1d, 3d, 7d, 10d, 21d, 28d | Medium to High expression | Low expression in Heart, High expression in Rear leg, Medium expression in rest |
| ASF1B | Amex3_2564976 | Low expression | Extremely high expression, decreases to medium expression at end | Medium expression in Rear leg, Low expression in rest |
| HELLS | Amex3_374305 | Low expression | Medium expression at stage 3, Low expression only at 1, 2,4,14,19,24 stages | No expression |
| HIST3H2BB | Amex3_161534 | High to Extremely high expression | Low expression only at 1,2,5,8,9,16 stage | No expression |
| HMGB2 | Amex3_403585 | No expression till 12hrs, then Low expression | Low expression till 8 stage, then Extremely high expression | Low to Medium expression |
| NASP | Amex3_2503343 | No expression till 6hrs, then Low expression | Extremely high expression | Medium expression in Liver, High expression in rest |
| RPA2 | Amex3_2565539 | Low expression | Extremely high to High expression | Low expression in Heart and Liver, Medium expression in rest |
| SART3 | Amex3_3258344 | Low expression | Extremely high expression | Medium expression |
| TOP1 | Amex3_1383108 | Low expression | Extremely high expression | Medium expression |
| TOP2B | Amex3_99453 | Low expression | Low expression till 7d, then medium expression till 10d, then Extremely high expression | High expression in Gills and Rear leg, Medium expression in rest |
| ACTA1 | Amex3_288061 | Medium expression at 3hrs,12hrs, Low expression in rest | Low expression till 21d, then High expression | Medium expression in Gills and Rear leg, Low expression in Heart |
| ACTN3 | Amex3_2626973 | No expression | Low expression at 8,9,24,40 stage | Extremely high expression in Tail, High expression in Front and Rear leg, Low expression in Heart |
| DES | Amex3_334965 | Medium expression till 1d, then Low expression | Low expression only at 1,3,14,16,19,24,40 stages | Low expression in Liver, Extremely high expression in rest |
| MYBPC2 | Amex3_77332 | Medium expression till 12hrs, then Low expression, gradually decreasing trend | Low expression only at 1,5,14,16,24,40 stages | Low expression in Heart and Liver, Extremely high expression in rest |
| MYBPC3 | Amex3_68648 | Low expression only at 0hrs,12hrs,1d,3d,5d,14d, 21d,28d | Low expression only at 19 stage | Extremely high expression in Heart, Low expression in Front leg and Gills, no expression in rest |
| MYH2 | Amex3_1761 | Low expression | Low expression only at 40 stage | No expression in Heart or Liver, Medium expression in Gills, Extremely high expression in rest |
| MYH4 | Amex3_1761 | Low expression | Low expression only at 40 stage | No expression in Heart or Liver, Medium expression in Gills, Extremely high expression in rest |
| MYH7 | Amex3_105660 | Low expression | Low expression only after 7 stage | Low expression in Liver, Medium expression in Gills, High expression in Tail, Extremely high expression in rest |
| MYL1 | Amex3_2633364 | High expression till 12hrs, then Medium expression till 5d, then Low expression | Low expression | No expression in Liver, Low expression in Heart, High expression in Gills, Extremely high expression in rest |
| MYL2 | Amex3_22809 | Low expression at 7d, 10d, 14d, Medium expression for rest | Low expression | No expression in Liver, Low expression in Heart, Medium expression in Gills, Extremely high expression in rest |
| MYL3 | Amex3_329288 | Extremely high expression till 12hrs, then High expression till 5d, then Low expression | Low expression only at 1,14 stage, Medium expression at 40 stage | Medium expression in Liver, Extremely high expression in rest |
| MYL4 | Amex3_2639977 | Low expression | Low expression | No expression in Front leg and Gills, Low expression in Liver and Tail, Medium expression in Rear leg, Extremely high expression in Heart |
| TNNC1 | Amex3_339993 | High expression till 12hrs, then Medium expression till 3d, then Low expression | High expression at 40 stage, Medium expression at 24 stage, low expression for rest | Low expression in Liver, Extremely high expression in rest |
| TNNC2 | Amex3_361070 | Extremely high expression, decreases to Low expression | High expression at 1 stage, then Low expression for rest | Low expression in Liver, High expression in Heart, Extremely high expression in rest |
| TNNI1 | Amex3_115364 | Extremely high expression till 12hrs, then High expression till 3d, then Low expression for rest | Low expression only at 2,5,10,12,16,19,24,40 stage | No expression in Liver, Medium expression in Heart, High expression in Gills, Extremely high expression for rest |
| TNNI2 | Amex3_2638627 | Extremely high expression till 1d, then high expression till 5d, then Low expression for rest | Extremely high expression for 1 stage, then Low expression for rest | No expression in Liver, Low expression in Heart, Extremely high expression in rest |
| TNNT1 | Amex3_198319 | High expression till 3d, then Low expression | Low expression only at 2,9,10,11,12,14,16,19,24,40 stage | No expression in Liver, Low expression in Heart, Medium expression in Gills, Extremely high expression in rest |
| TNNT3 | Amex3_823025 | Extremely high expression till 3d, then Medium and Low expression | High expression at 1 stage, then Low expression for rest | No expression in Liver, Medium expression in Heart, Extremely high expression in rest |
| TPM1 | Amex3_190914 | Extremely high expression till 5d, then Low expression | Medium expression | Medium expression in Liver, Extremely high expression in rest |
| TPM3 | Amex3_345240 | Medium expression till 3d, then Low expression | Low expression only at 9,10,11,12,19,24,40 stage | No expression in Liver, Extremely high expression in rest |
| ABAT | Amex3_2629439 | Low expression | Low / Medium expression | Extremely high expression in Liver, Medium expression in Heart, Low expression in rest |
| ANXA6 | Amex3_42541 | Low expression | Low expression | High expression in Gills, Extremely high expression in rest |
| BIN1 | Amex3_656102 | Low expression only at 0hrs,10d,21d,28d | Medium / High expression | Low expression only in Tail, no expression in rest |
| MYH10 | Amex3_3530194 | Low expression | Extremely high / High expression | Extremely high expression in Heart, Medium / High expression in rest |
| MYOZ1 | Amex3_2633493 | Medium / High expression till 7d, then Low expression | Low expression | No expression in Heart or Liver, High expression in Gills, Extremely high expression in rest |
| GAPDH | Amex3_128063 | Low expression only at 12hrs,1d | Medium expression at 40 stage, Low expression 1,2,6 stage, No expression in other stages | No expression |
| GPD1 | Amex3_132015 | Low expression | Low expression | Medium expression in Gills, Extremely high expression in liver, High expression in rest |
| GPI | Amex3_3534586 | High expression at 6hrs and 1d, Low expression at 10d, Medium expression in rest | High expression in 2,24 stage, Extremely high expression in rest | Extremely high expression |
| MDH2 | Amex3_124638 | Low expression | Low expression | Extremely high expression |
| PFKM | Amex3_2632336 | Low expression | Low expression | No expression in Liver, Low expression in Gills, High expression in Heart and Tail, Extremely high expression in rest |
| PGK1 | Amex3_375656 | High expression at 6hrs, Low expression at 10d, Medium expression for rest | Extremely high / High expression | Extremely high expression |
| SLC25A12 | Amex3_349153 | Low expression | Low expression | Low expression in Liver, High expression in Gills and Tail, Extremely high expression in rest |
| TPI1 | Amex3_2565813 | Low expression only at 0hrs, 6hrs,12hrs,1d,3d,7d,28d | Extremely high expression at 1,7,9,10,11,12,40 stage, High expression in rest | Extremely high expression |
| PGM1 | Amex3_3531432 | Low expression | Low expression | Low expression |
| PRKAG2 | Amex3_604 | Low expression only at 6hrs,1d,10d,14d | Medium expression till 12 stage, then Low expression for rest | Low expression in Tail and Heart, Medium expression for rest |
| ATP1B2 | Amex3_110410 | Medium expression | Low expression till 8 stage, High expression at 9 stage, then Extremely high expression | Low expression in Liver, Medium expression in Heart, Extremely high expression in rest |
| GABARAPL2 | Amex3_3107823 | Low expression | High expression at 2,40 stage, Medium expression at 12,24 stage, Low expression at 14,16,19 stage, Extremely high expression for rest | High expression in Liver, Extremely high expression in rest |

Figure 5 displays the overall results of all the genes, which are listed at the top of the graph, of each other the three data retrievals/comparisons we made: the regeneration process, development process, and the protein concentration in major tissues/organs. The colors show the relative expression levels. We see that regeneration, as observed above, has most genes were expressed at a low or no expression. We also observe that in embryo and major tissues more genes are expressed at higher quantities.

## DISCUSSION

### Regeneration Genes

We were able to find that there are orthologues genes conserved between humans and Axolotl gene sequences. Considering the functions of many genes and the proteins they produce in both organisms, especially for major organs and tissues, it can be observed that many of the same functions need to be carried out by the organisms for survival. That implies that they utilize similar if not the same proteins adapted best for the organism.

The gene expression observed in their respective transcripts showed that genes were expressed in a variety of levels and each condition displayed gene expression in different quantities.

In regeneration, most genes had low expression while a few had high and extremely high expression, displaying how fewer proteins are required for the regeneration process, and most genes are expressed in moderately lower levels but still play instrumental roles in the regeneration process. The few genes expressed in higher numbers are probably expressed in much higher quantities to make up and are required in large quantities to aid in the regenerative process.

In the embryo/development, our observation concludes that more proteins are required for the developmental process and more genes are required in large numbers, and this could be explained because development takes longer and requires more resources to complete correctly, compared to regeneration, which only is re-developing one part of the body at the time.

Finally, gene expression in the major organs demonstrated the vast number of genes and proteins required for the Axolotl’s regular body functions. There was also a considerable number of genes that were expressed in lower and medium quantities, showing that some functions only require small quantities to complete the necessary processes.

### Conservation in Other Species

Figure 8 shows the evolutionary relationship of different species of the *p53* cancer suppressor gene. The numbers on the branches of the phylogenetic trees are bootstrapping values.

**Figure 8.**
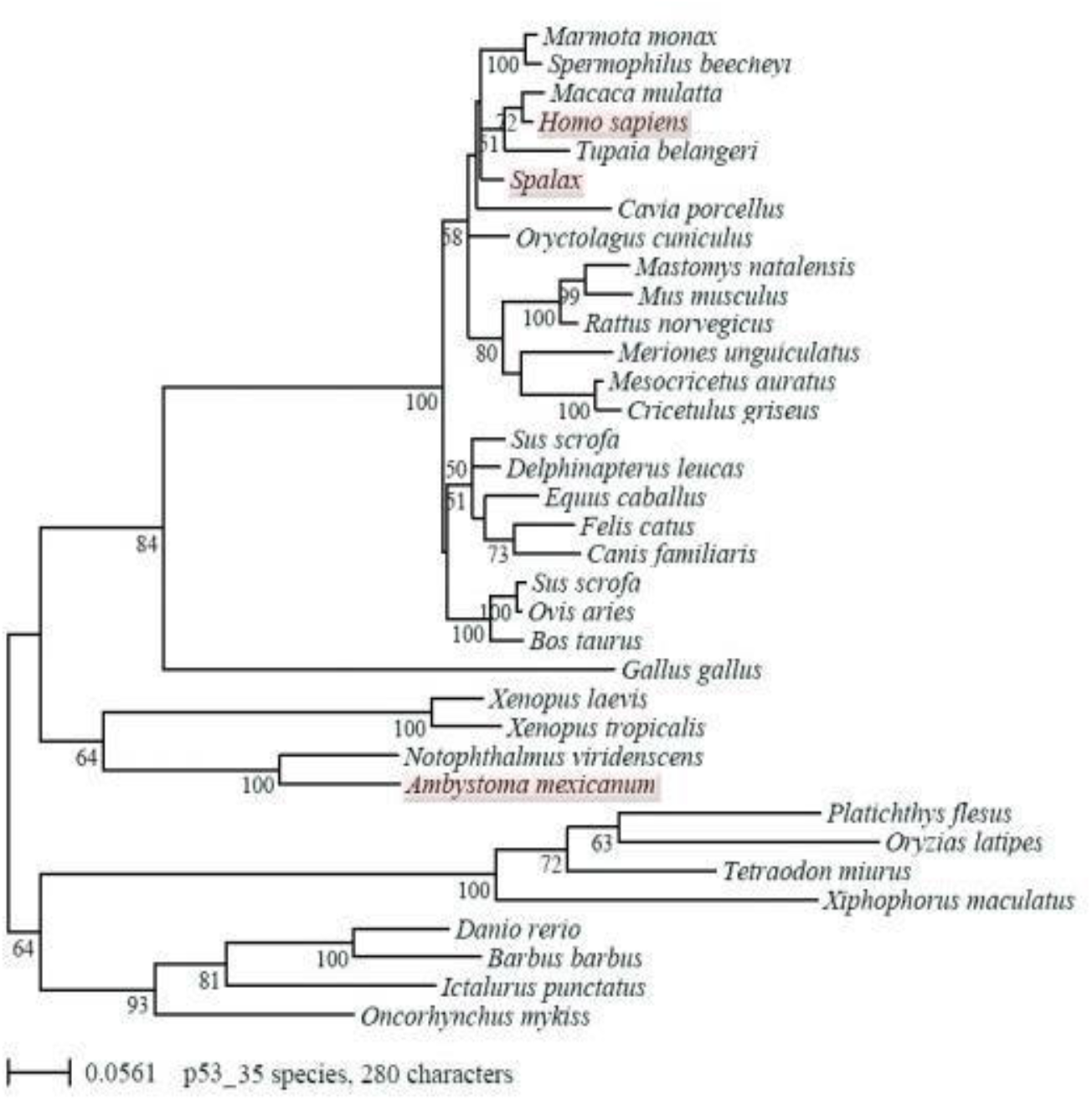
Phylogenetic analysis for the P53 gene (Villiard 2007).

Figure 9 is a phylogenetic tree of BH3-only proteins. Sequences found for zebrafish, Xenopus, human and axolotl were translated into amino acid sequences and aligned using ClustalW2 and T-coffee. A tree was generated and visualized. The scale bar indicates amino acid changes” (Vesna 2018). The Axolotl and Humans are shown to have evolutionary connections from the *Bad, Bid*, and *Bim* genes, where they are the closest relatives of those genes compared to the five organisms in the study.

**Figure 9.**
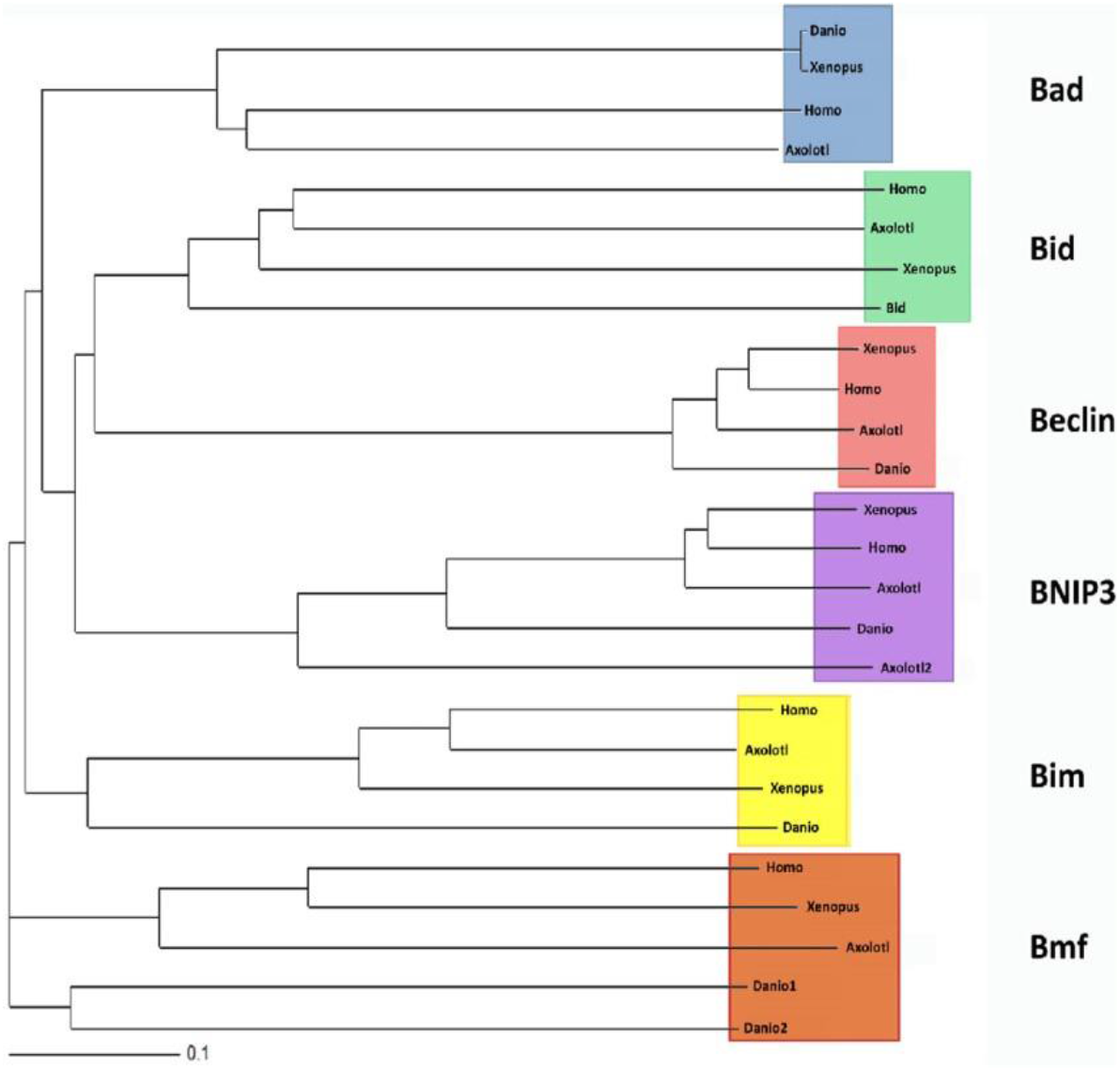
Phylogenetic analysis for BH3 protein family (Vesna 2018)

The Axolotl also shares characteristics with other species, as portrayed in both Figures. In Figure 9, the Eastern Newt (*Notophthalmus viridescens*) and Sub-Saharan Frogs (*Xenopus* sp) are most closely related with the Axolotl’s *p53* tumor suppressor gene. In Figure 9, the *Danio* species are most closely evolutionarily related to the Axolotl in the *Beclin* and *Bim* genes. Like this, as the Axolotl’s characteristics are more studied and their similarities are discovered with other organisms, more evolutionary relationships can be drawn and eventually the entire Axolotl genome will be linked with other organisms in a detailed and specific phylogenetic tree.

## CONCLUSION

The Axolotl genome is still relatively recent and there is still much to explore, we have only begun to scratch the surface of the regenerative capabilities Axolotls possess and future research is required to obtain more information on that ability.

The next steps for our research are to continue research into Axolotl’s and Human shared gene similarities and how genes can be manipulated to yield desirable results for regeneration.

For further research, topics can include: using comparisons for different human gene and finding the similarities between the human and Axolotl genome, studying the regenerative ability of other animals such as other salamanders as well, and uncovering the mysteries behind the other “superpowers” Axolotls have and utilizing them to help humans, such as Axolotl’s being more resistant to cancer.

## Funding

This study was supported by the SkoolMentor program [https://www.skoolmentor.com/].

## Acknowledgments

The authors acknowledge the Molecular and Developmental Complexity Group (Alfredo Cruz Lab) at UGA-LANGEBIO for generating and providing the data.

## Notes

### Competing Interest Statement

The authors have declared no competing interest.

### Summary of Updates

Harshil Shah's affiliation added, some minor errors fixed.

